# EquiPNAS: improved protein-nucleic acid binding site prediction using protein-language-model-informed equivariant deep graph neural networks

**DOI:** 10.1101/2023.09.14.557719

**Authors:** Rahmatullah Roche, Bernard Moussad, Md Hossain Shuvo, Sumit Tarafder, Debswapna Bhattacharya

## Abstract

Protein language models (pLMs) trained on a large corpus of protein sequences have shown unprecedented scalability and broad generalizability in a wide range of predictive modeling tasks, but their power has not yet been harnessed for predicting protein-nucleic acid binding sites, critical for characterizing the interactions between proteins and nucleic acids. Here we present EquiPNAS, a new pLM-informed E(3) equivariant deep graph neural network framework for improved protein-nucleic acid binding site prediction. By combining the strengths of pLM and symmetry-aware deep graph learning, EquiPNAS consistently outperforms the state-of-the-art methods for both protein-DNA and protein-RNA binding site prediction on multiple datasets across a diverse set of predictive modeling scenarios ranging from using experimental input to AlphaFold2 predictions. Our ablation study reveals that the pLM embeddings used in EquiPNAS are sufficiently powerful to dramatically reduce the dependence on the availability of evolutionary information without compromising on accuracy, and that the symmetry-aware nature of the E(3) equivariant graph-based neural architecture offers remarkable robustness and performance resilience. EquiPNAS is freely available at https://github.com/Bhattacharya-Lab/EquiPNAS.

## Introduction

Interaction of protein with Deoxyribonucleic acid (DNA) and Ribonucleic acid (RNA) underpins a wide range of cellular and evolutionary processes such as gene expression, regulation, and signal transduction ^1-4^. The identification of the interaction sites between proteins and nucleic acids (i.e., binding sites) is important for determining protein functions ^5^ and novel drug design ^6^. A number of computational methods for predicting protein-DNA and protein-RNA binding sites have been developed to overcome the challenges of lengthy and expensive nature of experimental characterization of protein-nucleic acid binding sites. Such computational methods can be broadly categorized into two categories: sequence-based and structure-aware methods. Sequence-based methods such as SVMnuc ^7^, NCBRPred ^8^, DNAPred ^9^, DNAgenie ^10^, RNABindRPlus ^11^, ConSurf ^12^, TargetDNA ^13^, SCRIBER ^3^, and TargetS ^14^ exploit readily available and abundant protein sequence information to predict binding sites. However, these methods lack structural information, which can limit their prediction accuracy. To overcome the challenge, structure-aware methods such as COACH-D ^15^, NucBind ^7^, DNABind ^16^, DeepSite ^17^, aaRNA ^18^, NucleicNet ^19^, GraphBind ^20^, and GraphSite ^21^ integrate available structural information for binding site prediction. While structure-aware methods usually achieve higher prediction accuracy than sequence-based methods, a vast majority of structure-aware methods rely on known structural information from the Protein Data Bank (PDB) ^22^ that are not as abundant as sequence information, limiting their large-scale applicability.

Promisingly, the recent breakthrough of AlphaFold2 ^23,24^ has enabled highly accurate prediction of single-chain protein structures from sequence information, providing new opportunities for replacing the experimentally solved structures with AlphaFold2-predicted structural models as input for binding site prediction at scale, without compromising on accuracy. While a recent protein-DNA binding site prediction method, GraphSite ^21^, has successfully used AlphaFold2-predicted protein structural models, effective utilization of predicted structures from AlphaFold2 for protein-RNA binding site prediction is yet to be explored. Alongside the AlphaFold2 breakthrough, a significant advancement has been made in pre-trained protein language models (pLM) ^25-30^ powered by attention-based transformers networks ^31^. pLMs have proven highly successful in various predictive modeling tasks including protein structure prediction ^28,30^, protein function prediction ^26,29^, and protein engineering ^27,32,33^. Despite their usefulness, the potential of pLMs in protein-DNA and protein-RNA binding site prediction tasks remains to be unlocked. Given the recent progress, a natural question arises: can we develop a generalizable computational framework that can harness the power of pLMs while leveraging the predicted structural information by AlphaFold2 for accurate prediction of protein-DNA and protein-RNA binding sites at scale?

Here, we present EquiPNAS, a new pLM-informed equivariant deep graph neural network framework for accurate protein-nucleic acid binding site prediction. EquiPNAS effectively leverages the pLM embeddings derived from the ESM-2 model ^30^ for improved protein-DNA and protein-RNA binding site prediction. The core of EquiPNAS consists of an E(3) equivariant graph neural network architecture ^34^, employing symmetry-aware graph convolutions that transform equivariantly with translation, rotation, and reflection in 3D space. Such an architecture has recently been shown to offer substantial accuracy gain while exhibiting remarkable robustness and performance resilience in our work on protein-protein interaction site prediction ^35^. Inspired by the notable successes of pLMs ^32,36-38^, here we integrate pLM embeddings from the encoder-only transformer architecture of ESM-2 to refine our sequence-based node features using the E(3) equivariant graph-based framework. By doing this, we are able to significantly reduce the dependence on the availability of evolutionary information which is not always abundant such as with orphan proteins or rapidly evolving proteins, thus enabling us to build generalizable and scalable models. In addition, our translation-, rotation-, and reflection-equivariant deep graph learning architecture provides richer representations for molecular data compared to invariant convolutions, offering robustness for graph structured data and particularly suitable when predicted protein structures are used as input ^35^.

Our method, EquiPNAS, consistently outperforms the state-of-the-art methods in several widely used benchmarking datasets for both protein-DNA and protein-RNA binding site prediction tasks. EquiPNAS exhibits remarkable robustness with only a minor performance decline when switching from experimental structures to AlphaFold2 predicted structural models as input, enabling accurate prediction of protein-DNA and protein-RNA binding sites at scale. The pLM embeddings used in EquiPNAS are sufficiently powerful that can dramatically reduce the dependence on the availability of evolutionary information, leading to a generalizable framework. In addition, the symmetry-aware nature of the E(3) equivariant graph-based neural architecture of EquiPNAS offers remarkable robustness and performance resilience, as verified directly through our ablation study. An open-source software implementation of EquiPNAS, licensed under the GNU General Public License v3, is freely available at https://github.com/Bhattacharya-Lab/EquiPNAS.

## Materials and methods

**Fig 1.**
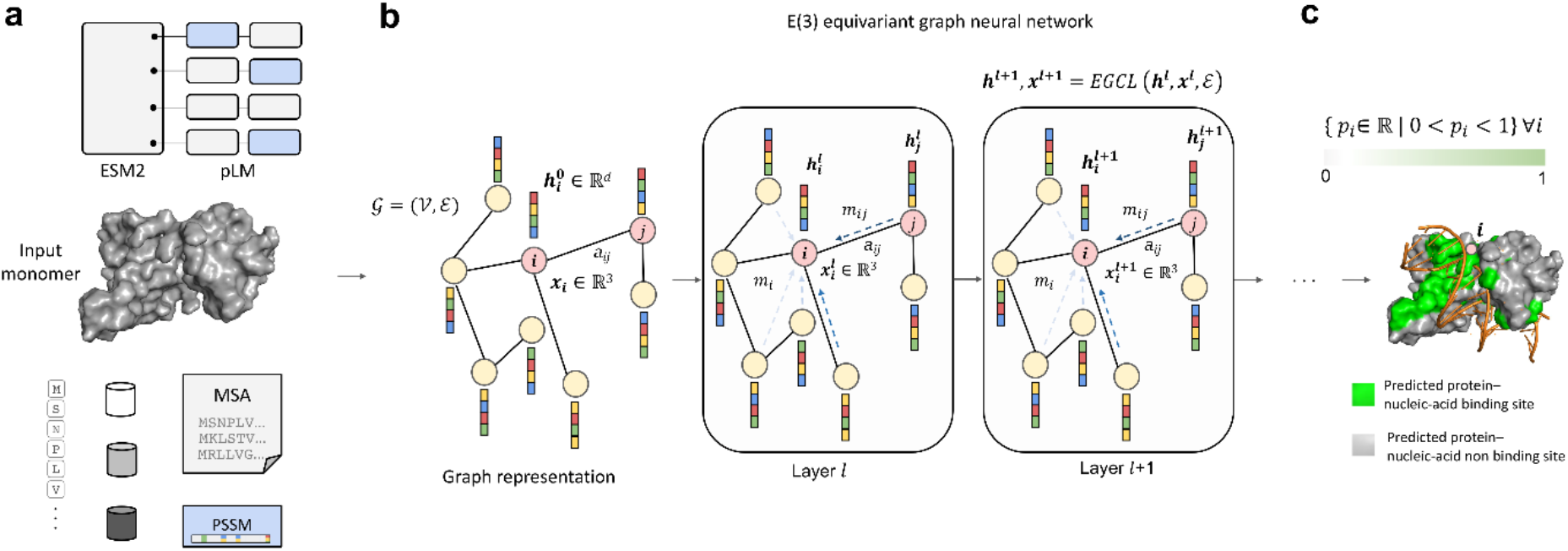
Illustration of the EquiPNAS method for protein-nucleic acid binding site prediction. (**a**) A set of node and edge features are generated from the input protein monomer. (**b**) E(3)-equivariant graph convolutions are employed on the featurized graph representation of the input. (**c**) Graph node classification is performed for residue-level binding site prediction.

### Overview of EquiPNAS Framework

**Fig. 1** illustrates our EquiPNAS method for protein-nucleic acid binding site prediction consisting of graph representation and featurization, E(3) equivariant graph neural network leveraging the coordinate information extracted from the input monomer together with sequence- and structure-based node and edge features as well as pLM embeddings from the ESM-2 model, and performing graph node classification to predict the probability of every residue in the input monomer to be a protein-nucleic acid binding site.

### Graph Representation and Feature Generation

#### Input Protein Graph Representation

We represent the input protein monomer as a graph 𝒢= (*𝒱, ε*), where each node *v ∈𝒱* represents a residue, and each edge *e ∈ε* represents an interacting residue pair. We consider a residue pair to be interacting if their Cα-Cα Euclidean distance is within 14Å for protein-DNA binding site prediction and 15Å for protein-RNA binding site prediction. The specific distance cut-offs are chosen through independent cross-validations for the protein-DNA and protein-RNA binding site tasks (**Supplementary Table 1, 2**). We additionally use a minimum sequence separation of 6 for the interacting residue pairs to focus on longer-range interactions.

**Table 1.** Sequence-based node features. The shape of the corresponding type for a protein with L residues is shown next to each feature.

| Features [*shape*] | Description |
| --- | --- |
| aa [*L*, 20] | One-hot encodings of 20 amino acid residue types. |
| PSSM [*L*, 20] | Normalized position specific scoring matrix (PSSM). |
| MSA [*L*, 256] | Multiple sequence alignment (MSA) representation distilled through ColabFold’s EvoFormer blocks. |
| pLM [*L*, 5120] | pLM embeddings from ESM-2 with 15B parameters. |

**Table 2.** Structures-based node features. The shape of the corresponding type for a protein with L residues is shown next to each feature.

| Features [*shape*] | Description |
| --- | --- |
| SS [*L, 11*] | One-hot encodings of 3- and 8-state secondary structure. |
| RSA [*L, 10*] | One-hot encodings of 2- and 8-state relevant solvent accessibility. |
| Local geometry [*L, 11*] | Cosine angle between the C=O of consecutive residues, normalized values of virtual bond and torsion angles, and normalized peptide backbone torsion angles. |
| Residue orientation [*L, 9*] | Unit vectors pointing towards the directions of $C_\alpha^{(i+1)} - C_\alpha^i$, $C_\alpha^{(i-1)} - C_\alpha^i$ and $C_\beta^i - C_\alpha^i$. |
| Relative residue positioning [*L, 2*] | Two types of relative positional features for the $i^{\text{th}}$ residue: (1) inverse of $i$ representing the relative sequence position, and (2) inverse of the Euclidean distance of $C_\alpha$ atom from the centroid representing the relative spatial positioning. |
| Residue virtual surface area [*L, 1*] | Virtual surface area of the conceptual convex hull constructed by the atoms in a residue. |
| Contact count [*L, 1*] | The number of spatial neighbors of each residue. |

#### Feature Generation

We use a number of standard sequence-derived node features including amino acid residue type, position specific scoring matrix (PSSM), multiple sequence alignment (MSA), and combine them with protein language model-based features from ESM-2 pLM. Additionally, we extract structure-derived node features from the input protein monomer, using either the experimentally solved structure or AlphaFold2-predicted structural model, including secondary structure (SS), relative solvent accessibility (RSA), local geometry, residue orientations, relative residue positioning, residue virtual area, and contact count.

#### Sequence-based Node Features

An overview of sequence-based node features and the corresponding shape can be found in **Table 1**. We use one-hot encoding to represent each of the 20 amino acid residue types (aa) as a binary vector with 20 entries. We run PSI-BLAST ^39^ to obtain position specific scoring matrix (PSSM). We then extract the first 20 columns of the PSSM and normalize the values using the sigmoidal function. We additionally generate multiple sequence alignment (MSA) from the input amino acid sequence by running ColabFold ^40^ pipeline, which uses MMseq2 ^41^ for MSA generation. The generated MSA is then fed to the EvoFormer blocks of AlphaFold2 as implemented in the ColabFold pipeline, resulting in a distilled MSA representation encoded as a dictionary. We extract the first row of the distilled MSA representation (‘msa_first_row’ from the dictionary) to be used as our MSA feature. We also use protein language model-based features from the pretrained ESM-2 model, having 15B parameters ^30^. Specifically, we use the ‘representations’ embeddings as pLM features by supplying the amino acid sequence to the ESM-2 model.

#### Structure-based Node Features

Our structure-based node features and the corresponding shape can be found in **Table 2**. We use one-hot encoding to represent both 3-state and 8-state secondary structures (SS). Additionally, we use one-hot-encodings to represent both 2-state relative solvent accessibility (RSA) features using an RSA cut-off of 50 and finer-grained 8-state RSA features by discretizing the RSA value into 8 bins with the following ranges: 0-30, 30-60, 60-90, 90-120, 120-150, 150-180, 180-210, and >210. We also extract local geometric features directly from the input protein monomer. These include the cosine angle between the C=O of consecutive residues, normalized virtual bond and torsion angles formed between consecutive Cα atoms, and normalized backbone torsion angles of the polypeptide chain. Inspired by the recent GVP-GNN study ^42^, we adopt two types of residue orientation features in our study: (1) unit vectors pointing towards Cα^(i+1)^ *−* Cα^i^ and Cα^(i*−*1)^ *−* Cα^i^, and (2) unit vectors indicating the imputed direction of Cβ^i^ *−* Cα^i^, which is computed assuming tetrahedral geometries and normalization. We use two types of relative residue positioning features for the i^th^ residue of the input protein monomer: (1) the relative sequence position captured by the inverse of i, and (2) the relative spatial positioning captured by the inverse of the Euclidean distance between the centroid of the input protein monomer and the Cα atom of the ith residue. We additionally conceptualize an amino acid residue as a virtual convex hull that is constructed by its constituent atoms and quantify the virtual surface area of the convex hull and calculate its inverse to use as a feature. Finally, we include the normalized contact count as a structure-driven feature, defined as the number of spatial neighbors of each residue (i.e., residues that are in contact) where two residues are considered to be in contact if the Euclidean distance between their Cβ atoms is less than 8Å.

#### Edge Features

As the edge feature for the graph 𝒢= (*𝒱, ε*), we use the ratio of the logarithm of the absolute difference between the indices of the two residues (*log* |*i-j*|) in the primary sequence and their Euclidean distance. The numerator of the ratio measures how far apart the two residues are in the primary sequence, while the denominator measures their spatial distance in 3D space.

#### Coordinate Features

We obtain coordinate features from the Euclidean coordinates (x, y, and z) of the Cα atoms in input protein monomers.

### Network Architecture

Our network architecture consists of deep E(3)-equivariant graph neural networks (EGNNs), independently trained for protein-DNA and protein-RNA binding site prediction tasks. The input to the EGNNs includes the node and edge features described above as well as coordinate features based on the Cartesian coordinates of the Cα atoms in the input protein monomer. The EGNN architecture consists of a stack of equivariant graph convolution layers (EGCL), performing a series of transformations of its input by updating the coordinate and node embeddings using the edge information and the coordinate and node embeddings from the previous layer. A linear transformation is first applied to the input node features 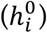, which results in a transformed set of node embeddings 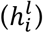. These embeddings, along with input coordinates 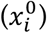 and edge information (*aij*) are passed to the subsequent EGCL layers. The EGCL operation attains equivariance by changing the standard message passing to equivariant message passing and by introducing coordinate updates. Unlike off-the-shelf graph neural networks that aggregate messages only from the neighboring nodes, equivariant graph neural networks aggregate messages from the whole graph. Additionally, an attention embedding is employed through a linear transformation on the aggregated message embedding, followed by a sigmoidal non-linear transformation. Finally, a linear transformation is applied to squeeze the hidden dimension of the last EGCL for condensing the learned information into a single scalar value, followed by a sigmoidal function to obtain the node-level classification to predict the likelihood of every residue in the input monomer to be a protein-nucleic acid binding site. The architecture of our EGNN consists of 12 EGCL layers with hidden dimensions of 768. The size of hidden dimensions and the number of layers are selected through 5-fold cross-validation (see **Supplementary Table 1, 2**). To mitigate the risk of overfitting, we apply dropout regularization to the node embeddings of each EGCL layer with a dropout rate of 0.1, determined through 5-fold cross validation (see **Supplementary Table 1, 2**). Our EquiPNAS models are implemented using PyTorch 1.12.0 ^43^ and the Deep Graph Library (DGL) 0.9.0 ^44^. During training, we use the binary cross-entropy loss function and a cosine annealing scheduler from the Stochastic Gradient Descent with Warm Restarts (SGDR) algorithm ^45^. We also utilize the ADAM optimizer ^46^, with a learning rate of 1e-4 and a weight decay of 1e-16. The training process consists of at most 40 epochs on an NVIDIA A40 GPU. In addition to the full-fledged version of EquiPNAS, we train baseline models for both protein-DNA and protein-RNA binding site prediction using the same hyperparameters and features as EquiPNAS, but without equivariant updates, that is, invariant baseline networks with the coordinate updates of the equivariant graph convolution layers turned off, enabling us to verify the importance of equivariance used in our model.

### Datasets and Performance Evaluation

For a fair performance comparison of our method against the state-of-the-art methods for protein-DNA and protein-RNA binding site prediction, we use widely recognized public datasets as follows.

#### Protein-DNA Benchmarking Dataset

To evaluate the performance of protein-DNA binding site prediction method, we use train (Train_573) and test (Test_129) datasets from the published work of GraphBind ^20^, which contain a total of 573 and 129 protein chains, respectively. Additionally, we use another test set consisting of 181 protein chains (Test_181) from the published work of GraphSite ^21^. These datasets are originally curated from the public BioLiP database ^47^ that contains precomputed protein-DNA and protein-RNA binding sites from known protein-DNA and protein-RNA complexes after filtering out protein chains with > 30% sequence similarity among the datasets, by applying CD-Hit ^48^ to ensure non-redundancy. The training dataset (Train_573) was released before 6 January 2016 whereas the Test_129 set was released between 6 January 2016 to 5 December 2018, and Test_181 was more recently released between 6 December 2018 to August 2021. The binding (and non-binding) residue count for Train_573, Test_129, and Test_181 are 14,479 (and 145,404), 2,240 (and 35,275), and 3,208 (and 72,050), respectively.

#### Protein-RNA Benchmarking Dataset

To evaluate the performance of protein-RNA binding site prediction method, we use the Train_495 set for training and the Test_117 set for testing, also from the published work of GraphBind ^20^, which contain a total of 495 and 117 protein chains, respectively. These datasets are also extracted from the BioLiP database ^47^ and pre-processed to ensure non-redundancy between the train and test sets, using CD-Hit ^48^ to filter out protein chains with >30% sequence similarity. The Train_495 set contains 14,609, and 122,290 binding, and non-binding residues, respectively, while in the Test_117 set, 2,031, and 35,314 residues are binding, and non-binding residues, respectively.

#### Evaluation metrics and competing methods

We assess the performance of our method using two widely recognized metrics: the area under the Receiver Operating Characteristic curve (ROC-AUC) and the area under the Precision-Recall curve (PR-AUC) scores. Both ROC-AUC and PR-AUC are threshold-independent metrics, thereby providing a comprehensive and robust view of the performance of a model across the full range of possible classification thresholds.

We compare our protein-DNA interaction site prediction method against eight existing methods. Three of the methods, SVMnuc ^7^, NCBRPred ^8^, and DNAPred ^9^, are sequence-based methods, while the other five methods, COACH-D ^15^, NucBind ^7^, DNABind ^16^, GraphBind ^20^, and GraphSite ^21^ are structure-aware methods. SVMnuc is a support vector machine (SVM)-based method that utilizes features from PSI-BLAST ^39^, PSIPRED ^49^, and HHblits ^50^. NCBRPred employs bidirectional Gated Recurrent Units (BiGRU) ^51^ with multi-label sequence labeling. DNAPred is a two-stage ensembled hyperplane-distance-based support vector machine (E-HDSVM) ^9^ for predicting protein-DNA binding sites. COACH-D is a consensus-based approach incorporating four different template-based and one template-free prediction methods. NucBind integrates the *ab initio* SVMnuc and template-based COACH-D for higher accuracy prediction. DNABind is a hybrid method combining machine learning with template-based predictions. GraphBind proposes hierarchical graph neural networks, while GraphSite employs graph transformer neural networks. Among these competing methods, GraphBind and GraphSite are the most recent and represent the state-of-the-art for protein-DNA binding site prediction.

We compare our protein-RNA binding site prediction method with seven existing methods. Two of the methods RNABindRPlus ^11^ and SVMnuc ^7^ are sequence-based methods, while the other five methods, COACH-D ^15^, NucBind ^7^, aaRNA ^18^, NucleicNet ^19^, and GraphBind ^20^ are structure-aware methods. SVMnuc, COACH-D, NucBind, and GraphBind are the methods we also compared against on protein-DNA binding tasks, as discussed earlier. RNABindRPlus is a hybrid method that combines sequence-homologs and support vector machine (SVM)-based predictions. aaRNA is a both sequence- and structure-based method that utilizes homology modeling to extract structural features along with various sequence-based features. NucleicNet is a deep learning framework that extracts physiochemical characteristics of the protein surface by quantifying it with grid points. Among these methods, GraphBind is currently the top-performing method for protein-RNA binding site prediction.

## Results

### Test Set Performance

**Table 3** shows the performance of EquiPNAS for protein-DNA (on Test_129 and Test_181 sets) and protein-RNA (on Test_117) binding site prediction tasks using AlphaFold2 predicted structural models as input compared to two closest competing methods: hierarchical graph neural network-based method GraphBind for protein-DNA and protein-RNA binding site prediction ^20^ and graph transformer-based method GraphSite for protein-DNA binding site prediction ^21^ (see **Supplementary Table 3** and **Supplementary Table 4** for comprehensive performance comparison against all competing methods). The results demonstrate that EquiPNAS attains the highest scores in all three test datasets. The performance gain of EquiPNAS over the state-of-the-art methods is particularly noteworthy considering PR-AUC, a stringent and rigorous evaluation metric. For example, EquiPNAS yields 56.9% relative PR-AUC gain over GraphBind for protein-RNA binding site prediction; and 14.5%-21.1% relative PR-AUC gains over GraphBind and 4.1%-4.6% relative PR-AUC gains over GraphSite for protein-DNA binding site prediction. In summary, EquiPPIS improves upon the state-of-the-art accuracy of both protein-DNA and protein-RNA binding site prediction using AlphaFold2 predicted structural models by consistently attaining better performance than the existing approaches.

**Table 3.**
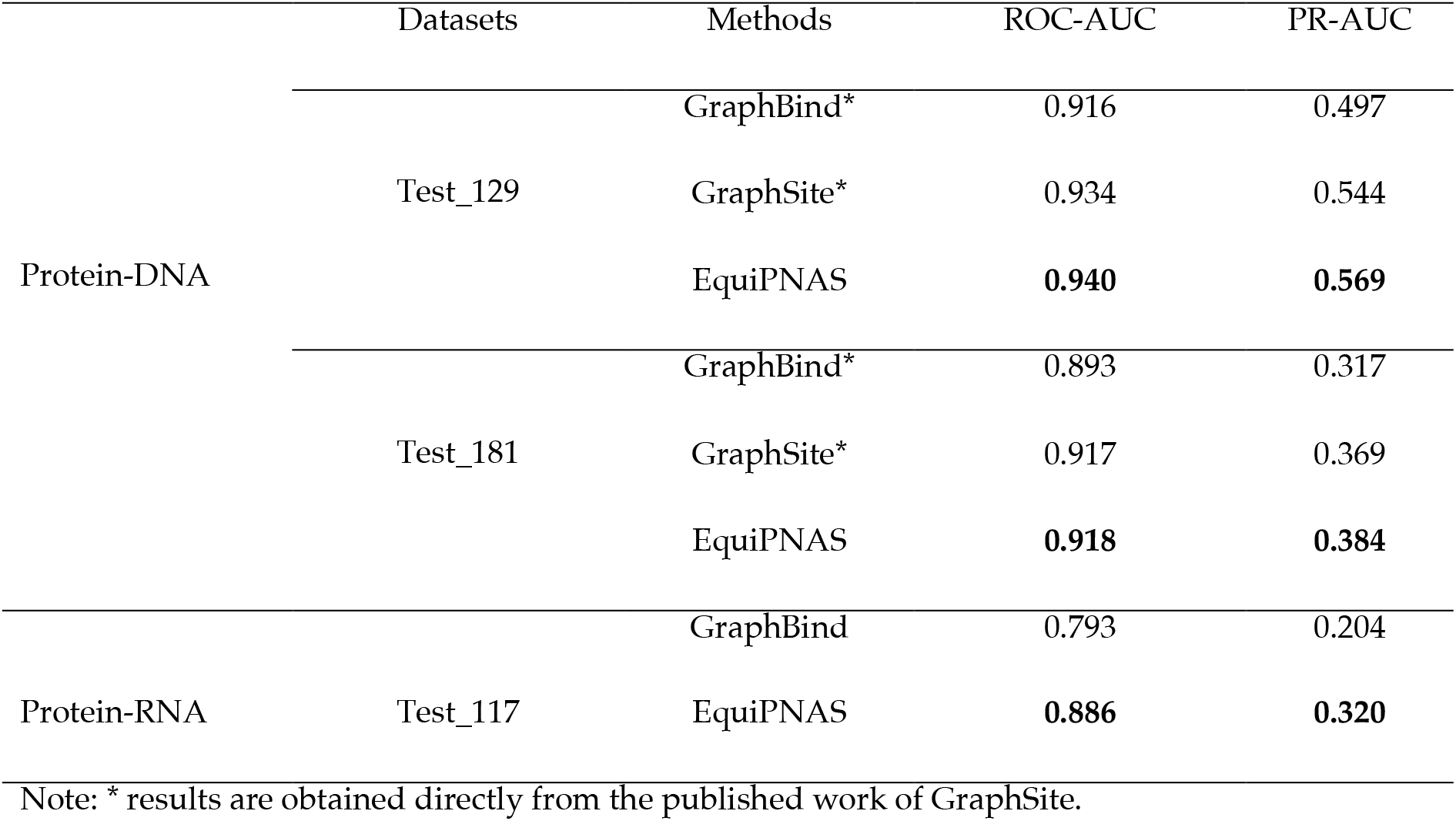
Protein-DNA and protein-RNA binding site prediction performance of EquiPNAS against the top-performing methods on the test datasets using AlphaFold2 predicted structural models as input. Values in bold represent the best performance.

**Table 4.** Protein-DNA and protein-RNA binding site prediction performance of EquiPNAS variant trained without any evolutionary information (w/o MSA + PSSM) against the top-performing methods on the test datasets using AlphaFold2 predicted structural models as input. Values in bold represent the best performance.

|  | Datasets | Methods | ROC-AUC | PR-AUC |
| --- | --- | --- | --- | --- |
| Protein-DNA | Test_129 | GraphBind* | 0.916 | 0.497 |
|  |  | GraphSite* | 0.934 | **0.544** |
|  |  | EquiPNAS w/o (MSA+PSSM) | **0.936** | **0.544** |
|  | Test_181 | GraphBind* | 0.893 | 0.317 |
|  |  | GraphSite* | **0.917** | **0.369** |
|  |  | EquiPNAS w/o (MSA+PSSM) | **0.917** | 0.364 |
| Protein-RNA | Test_117 | GraphBind | 0.793 | 0.204 |
|  |  | EquiPNAS w/o (MSA+PSSM) | **0.877** | **0.299** |

Note: \* results are obtained directly from the published work of GraphSite.

**Fig 2** presents 12 representative examples from the test datasets comparing the protein-DNA and protein-RNA binding site predictions using EquiPNAS against the second-best predictors: 4 from protein-DNA Test_129 (**Fig 2 a**), 4 from protein-DNA Test_181 (**Fig 2b**), and 4 from protein-RNA Test_117 (**Fig 2c**). The first two examples represent two human protein-DNA interactions: Transcription of Homo sapiens, Mus musculus (PDB ID: 5nj8, chain A), and Hydrolase/DNA of Homo sapiens, DNA launch vector pDE-GFP2 (PDB ID: 5t4i, chain B) as shown in **Fig 2a**. GraphSite fails to predict the vast majority of protein-DNA binding sites as reflected in its low F1-score, Matthew’s Correlation Coefficient (MCC) and PR-AUC in these two targets. In contrast, EquiPNAS achieves reasonably accurate prediction, with a remarkable gain of 0.506 and 0.417 points in F1-score, 0.458 and 0.357 points in MCC, and 0.381 and 0.381 points in PR-AUC, respectively. The third and the fourth examples, Splicing of Caenorhabditis elegans, synthetic construct (PDB ID: 5tkz, chain A) and Transcription of Escherichia coli (PDB ID: 5ond, chain A), both show inaccurate binding site prediction by GraphSite, resulting in predicting 5 (out of total 89 residues), and 25 (out of total 135 residues) false positives, respectively, which are noticeably high. EquiPNAS accurately predicts these binding sites, with only 1 (out of total 89 residues) and 7 (out of total 135 residues) false positives, respectively. GraphSite also generates inaccurate predictions for DNA binding protein/DNA in Escherichia coli (PDB ID: 6nua, chain A) and for Mycolicibacterium smegmatis MC2 155, Mycolicibacterium smegmatis (PDB ID: 6iod, chain A), with 28 (out of total 227 residues), and 32 (out of total 212 residues) false positives, respectively; whereas EquiPNAS achieves a much better overall prediction performance with only 3 (out of total 227 residues) and 9 (out of total 212 residues) false positives, respectively. Interestingly, EquiPNAS attains perfect prediction with both ROC-AUC and PR-AUC values of 1.0, as well as an F1-score and MCC of approximately 0.93 for a smaller target (73 residues), a transcription protein in Mycobacterium tuberculosis (PDB ID: 7kuf chain A). In contrast, GraphSite’s prediction is contaminated by several false positives, resulting in F1-score and MCC values of less than 0.65. Additionally, for an RNA binding protein/DNA in Homo sapiens (PDB ID: 7csz, chain A), our method still outperforms GraphSite, with a performance gain of 0.27 points in PR-AUC, 0.187 points in F1-score, and 0.216 points in MCC, whereas GraphSite fails to identify majority of binding site residues,particularly for DNA chain C, resulting in a high number of false negatives. The RNA binding protein example in Danio rerio and Caenorhabditis elegans (PDB ID: 6fq3, chain A) provides a remarkable demonstration of the superior performance of EquiPNAS in predicting protein-RNA binding sites, as compared to the closest competing method GraphBind. While GraphBind fails to accurately detect any binding site, with PR-AUC, F1-score, and MCC of 0.024, 0, and -0.017, respectively, EquiPNAS performs reasonably accurate predictions with much better PR-AUC, F1-score, and MCC of 0.732, 0.545, and 0.555, respectively. Furthermore, EquiPNAS shows highly accurate prediction for the transcription factor in Saccharomyces cerevisiae (PDB ID: 5o1y, chain A), exceeding GraphBind by 0.351 points in F1-score, 0.382 points in MCC, and 0.303 points in PR-AUC. Additionally, in comparison to EquiPNAS, GraphBind exhibits suboptimal performance due to both false positive and false negative predictions for the binding sites of OXIDOREDUCTASE/RNA in Escherichia coli (PDB ID: 5hr7, chain B) and Hydrolase/RNA Methanococcus maripaludis C5, Methanococcus maripaludis (PDB ID: 4z7l, chain A).

**Fig 2.**
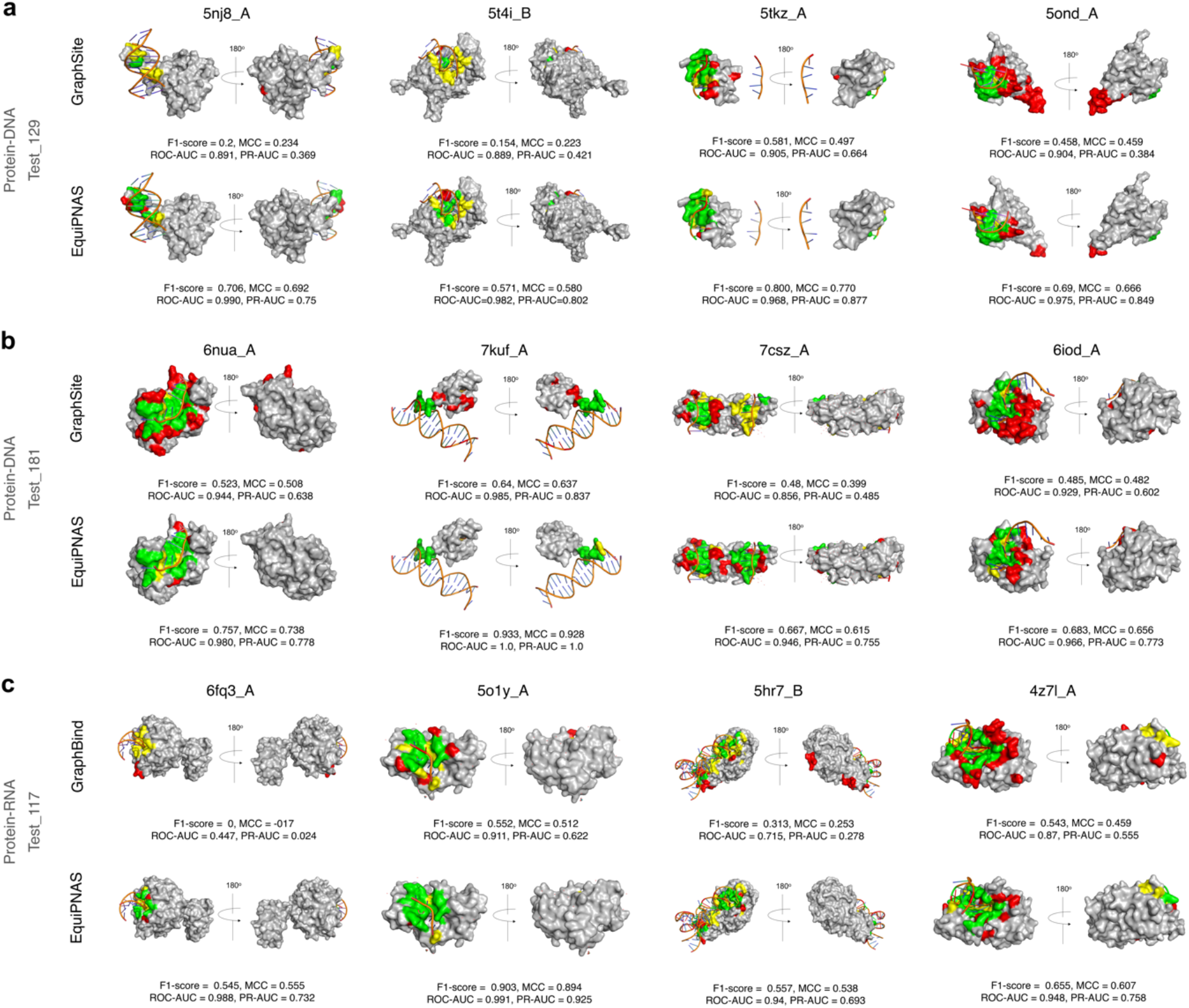
Representative examples of protein-DNA and protein-RNA binding site predictions using EquiPNAS and the closest competing methods compared to the experimental observation. For targets from the Test_129 (**a**) and Test_181 (**b**) sets, protein-DNA binding site prediction using GraphSite vs. EquiPNAS are shown. For targets from the Test_117 set (**c**), protein-RNA binding site prediction using GraphBind vs. EquiPNAS are shown. True Positive (TP), False Positive (FP), and False Negative (FN) binding sites are represented in green, red, and yellow, respectively.

In the above experiments, all methods use AlphaFold2 predicted structural models as input with EquiPNAS consistently delivering improved performance for both protein-DNA and protein-RNA binding site prediction tasks. However, structure-aware protein-nucleic acid binding site prediction methods traditionally rely on experimentally solved structures as input. Intuitively, using experimental structures as input, whenever available, should lead to better performance than using predicted structural models as input. Consequently, a natural question to ask is: How much performance decline do these methods suffer from when switching from experimental input to prediction? Not surprisingly, as shown in **Supplementary Table 3** and **Supplementary Table 4**, using experimental input leads to better accuracy in almost all cases. Promisingly, the performance decline of EquiPNAS when switching from experimental input to AlphaFold2 prediction is much smaller compared to other methods. For instance, EquiPNAS loses only ∼2.3% of PR-AUC points when using AlphaFold2 predictions as input instead of experimental ones for protein-DNA binding site prediction, whereas GraphBind experiences a higher PR-AUC drop of 4.4%-6.9% PR-AUC points. EquiPNAS also demonstrates robustness in protein-RNA binding site prediction with a negligible drop in ROC-AUC (0.1%) when using AlphaFold2 predictions as input, whereas GraphBind shows a much higher ROC-AUC drop (7.7%). That is, EquiPNAS exhibits a minor performance decline when switching from experimental input to prediction while outperforming both GraphBind and GraphSite regardless of the use of predicted or experimental structures, demonstrating its robustness and generalizability and enabling accurate prediction of protein-DNA and protein-RNA binding sites at scale using AlphaFold2 predicted structural models.

In the context of large-scale protein-nucleic acid binding site prediction using AlphaFold2 predicted structural models, a related question is: Is there any relationship between the self-estimated accuracy of AlphaFold2 predicted structural models and the accuracy of EquiPNAS binding site prediction? We examine the self-estimated accuracy of AlphaFold2 predicted structural models using the AlphaFold2 predicted local distance difference test (pLDDT) and the ROC-AUC and PR-AUC of EquiPNAS binding site prediction resulting from the predicted structure. Using a pLDDT threshold of 0.85, we divide the targets in the test sets into two roughly equal groups: moderate confidence predictions with pLDDT values *≤* 0.85 and high confidence predictions with pLDDT values > 0.85. **Supplementary Fig 1** shows the ROC-AUC and PR-AUC distributions for the two groups. Across the test datasets, high confidence predictions lead to better ROC-AUC and PR-AUC values compared to moderate confidence predictions, with the ROC-AUC and PR-AUC distributions resulting from the high confidence predictions skewed towards higher accuracy binding site prediction. Furthermore, we observe a noticeable difference in binding site prediction accuracy in terms of mean ROC-AUC and PR-AUC values resulting from the moderate confidence predictions versus the high confidence predictions (see **Supplementary Table 5**), indicating that the self-estimated accuracy of AlphaFold2 predicted structural models can inform the accuracy of EquiPNAS binding site prediction in the absence of any experimental information in that highly confident AlphaFold2 predictions tend to yield more accurate binding site prediction

### Ablation Study

#### Contribution of the pLM embeddings

EquiPNAS utilizes pLM embeddings from the pretrained ESM-2 model ^30^ as part of the sequence-based features. To evaluate the relative contribution of the protein language model-based features compared to the evolutionary features such as PSSM and MSA, we conduct a feature ablation study by excluding protein language model-based features or the evolutionary features from the full-fledged EquiPNAS feature set. **Fig 3** displays the 5-fold cross-validation performance of the ablated variants of EquiPNAS in terms of ROC-AUC and PR-AUC values for protein-DNA and protein-RNA binding site prediction. The results demonstrate that excluding pretrained protein language model-based features (no pLM) results in the worst performance with a relative PR-AUC drop of 18.5% (**Fig 3b**) and 15.4% (**Fig 3d**) for protein-DNA and protein-RNA binding site predictions, respectively. Such a significant performance drop highlights the importance of using pLM embeddings for our prediction. In contrast, we observe only minor performance drops when one or both evolutionary features were discarded. Even discarding both the evolutionary features (No (PSSM+MSA)) results in a relative PR-AUC drop of only 2.8% and 2% for protein-DNA and protein-RNA binding site predictions, respectively. Overall, compared to the relatively minor but positive contribution of evolutionary features, protein language model-based features have a major contribution to the improved performance of the new EquiPNAS model.

**Fig 3.**
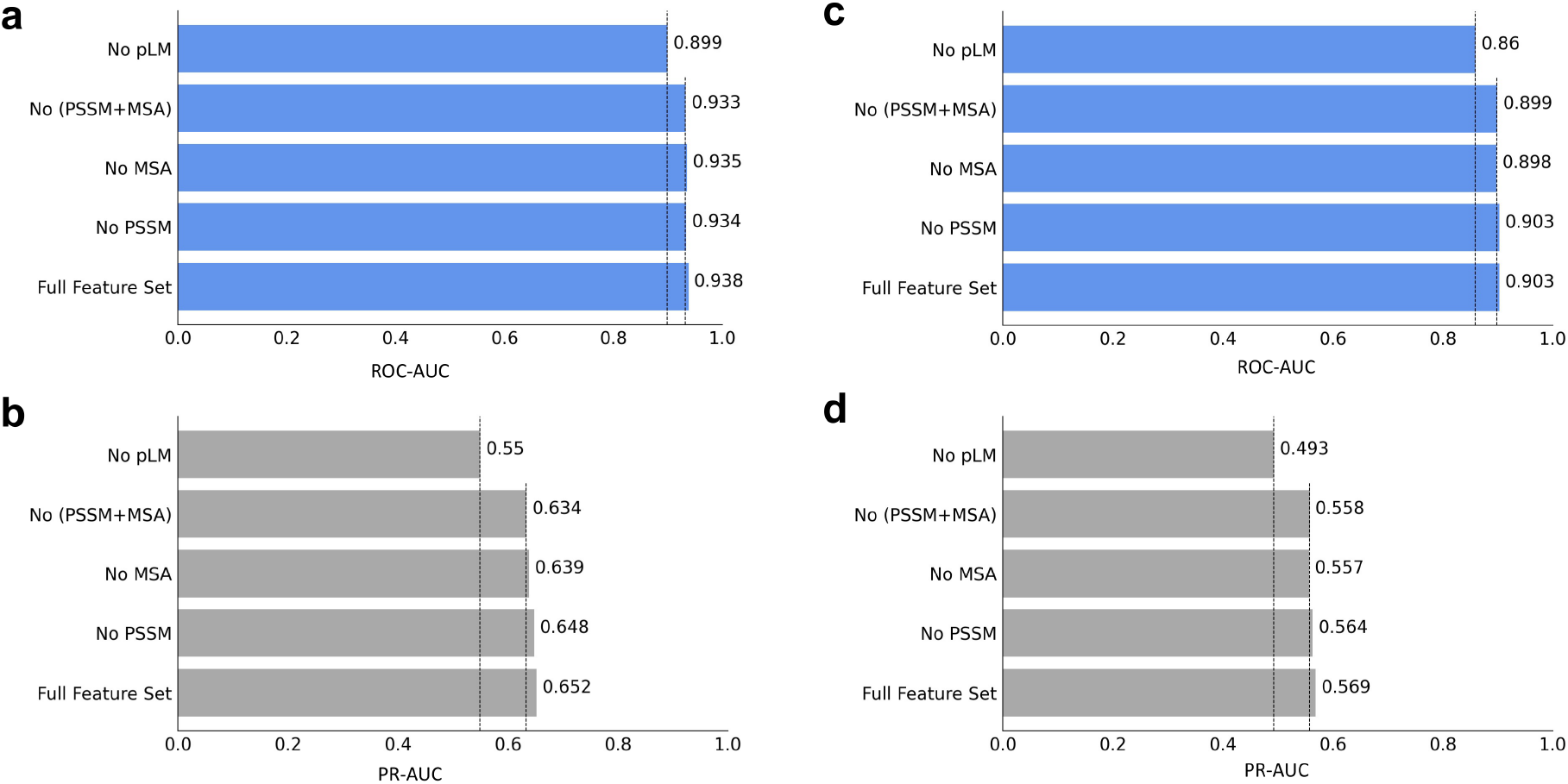
Feature ablation study. For protein-DNA binding site prediction, bar charts representing the performance of the ablated variants in terms of (**a**) ROC-AUC and (**b**) PR-AUC obtained using 5-fold cross validation are shown. For protein-RNA binding site prediction, bar charts representing the performance of the ablated variants in terms of (**c**) ROC-AUC and (**d**) PR-AUC obtained using 5-fold cross validation are shown.

Recognizing the major contribution of pLM features compared to the relatively minor impact of the evolutionary features, we investigate the performance of our method utilizing the pLM embeddings, but without using any evolutionary information. Specifically, we discard the PSSM and MSA features and retrain our method on the full training set, and evaluate the performance on the test sets for both protein-DNA and protein-RNA binding site prediction tasks. As reported in **Table 4**, We find that for protein-DNA binding site prediction, EquiPNAS without PSSM or MSA (denoted by “EquiPNAS w/o (PSSM+MSA)”) outperforms GraphBind, and performs comparably to GraphSite; with only a slight performance decline compared to the full-fledged version of EquiPNAS. For example, in Test_129, EquiPNAS w/o (PSSM+MSA) achieves a ROC-AUC of 0.936 and a PR-AUC of 0.544, which is comparable to GraphSite (ROC-AUC of 0.934 and PR-AUC of 0.544) and much higher than GraphBind (ROC-AUC of 0.916 and PR-AUC of 0.497). We observed a similar trend in Test_181. In contrast, the state-of-the-art GraphSite experiences a noticeable performance drop without using any of evolutionary features. As reported in the published work of GraphSite, PR-AUC drops from 0.544 down to 0.452 without using its MSA-derived features (-AF2 Single). For protein-RNA binding site prediction (Test_117), EquiPNAS w/o (PSSM+MSA) achieves a ROC-AUC of 0.877 and a PR-AUC of 0.299, which is noticeably better than GraphBind (ROC-AUC of 0.793 and PR-AUC of 0.204). Collectively, the results demonstrate the robustness of EquiPNAS over the state-of-the-art methods in that EquiPNAS is able to significantly reduce the dependence on the availability of evolutionary information which is not always abundant such as with orphan proteins or rapidly evolving proteins. Even without using any evolutionary information, and thus at a much lower computational overhead required for MSA and PSSM feature generation, our method performs comparably (in the case of protein-DNA), even superior (in the case of protein-RNA) to the full-fledged state-of-the-art protein-DNA and protein-RNA binding site prediction methods. In summary, EquiPNAS enables to build generalizable and scalable models.

#### Contribution of equivariance

EquiPNAS delivers robust and improved performance across various datasets and predictive modeling scenarios. In order to understand the reasons behind such improved performance and verify that it is connected to the equivariant nature of the model, we perform an ablation study by isolating the effect of the equivariant graph convolutions used in EquiPNAS. In particular, we train a family of baseline graph neural networks for protein-DNA and protein-RNA binding site prediction tasks after turning off the coordinate updates of the equivariant graph convolution layers, thus making it an invariant network. Both the equivariant (the full-fledged version of EquiPNAS) and invariant counterparts are trained on the same training datasets using the same set of input features and hyperparameters as the full-fledged version of EquiPNAS. **Fig 4** shows the the performance of the equivariant and invariant networks using both experimentally determined (native) and AlphaFold2 predicted structures. The results demonstrate that equivariant networks used in the full-fledged version of EquiPNAS consistently outperform the invariant baseline networks regardless of the use of predicted or native structures as input. Strikingly, the invariant baseline models even using the native structures perform worse than the equivariant models using the AlphaFold2 predicted structures, let alone the equivariant models using the experimental structures. For instance, in the Test_129 set, the baseline invariant model attains ROC-AUC (and PR-AUC) of 0.938 (and 0.565) using the native structures, whereas the equivariant model attains ROC-AUC (and PR-AUC) of 0.940 (and 0.569) using AlphaFold2 predicted structures, and 0.943 (0.582) using native structures. A similar trend is also overserved in test sets Test_181 and Test_117. Overall, the results highlight the performance contribution and remarkable robustness of the equivariant networks used in EquiPNAS, attaining better accuracy with AlphaFold2 predicted structural models than what an invariant counterpart can achieve even with experimental structures for both protein-DNA and protein-RNA binding site prediction tasks.

**Fig 4.**
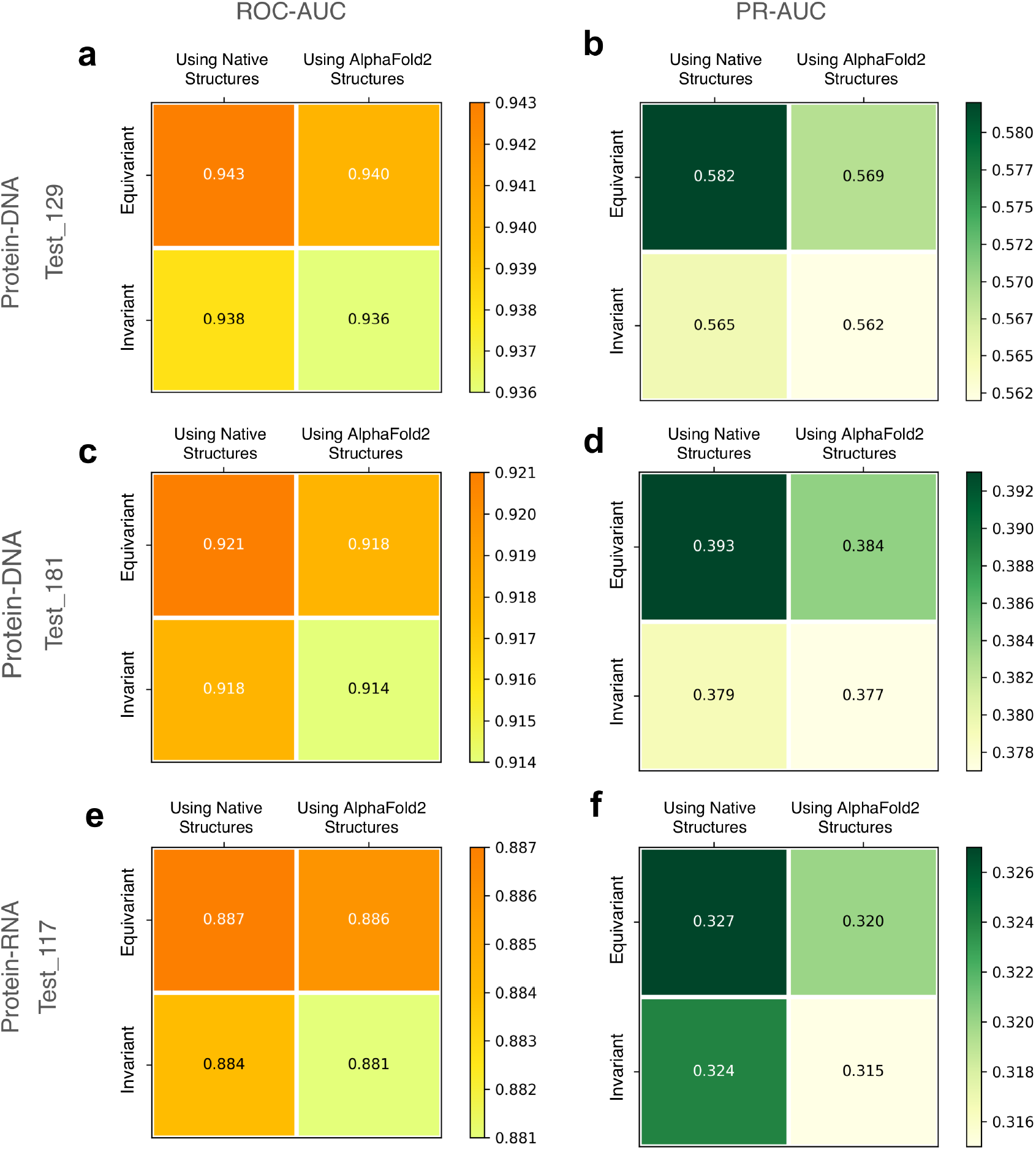
The performance of equivariant networks used in the full-fledged version of EquiPNAS compared against the invariant baseline networks using both experimental (native) and AlphaFold2 predicted structures as input. ROC-AUC and PR-AUC for protein-DNA test set Test_129 are presented in (**a, b**); ROC-AUC and PR-AUC for protein-DNA test set Test_181 are presented in (**c, d**); ROC-AUC and PR-AUC for protein-RNA test set Test_117 are presented in (**e, f**).

## Discussion

This work presents EquiPNAS, a new pLM-informed equivariant deep graph neural network framework for accurate protein-nucleic acid binding site prediction. We demonstrate that EquiPNAS consistently outperforms the state-of-the-art methods on both protein-DNA and protein-RNA binding site prediction tasks. A major contribution of our work is the successful utilization of protein language model (pLM) embeddings, a previously unexplored avenue in the context of protein-DNA and protein-RNA binding site predictions. Our ablation study reveals that the pLM embeddings are sufficiently powerful that can dramatically reduce the dependence on the availability of evolutionary information which is not always abundant such as with orphan proteins or rapidly evolving proteins, enabling us to build generalizable models. Moreover, despite being trained on experimental structures as input, our method exhibits remarkable robustness and performance resilience by attaining high predictive accuracy even when AlphaFold2 predicted structural models are used as input, dramatically enhancing the scalability of protein-nucleic acid binding site prediction without compromising on accuracy. Through controlled experiments, we directly verify that the symmetry-aware nature of the E(3) equivariant graph-based framework is a major driving force behind the improved performance of EquiPNAS, particularly when predicted structures are used as input.

While this work focuses on partner-independent protein-nucleic acid binding site prediction, that is, predicting the binding sites based only upon the surface of an isolated protein without any prior knowledge about the interacting nucleic acid partner; incorporating additional information regarding the DNA or RNA molecules interacting with the protein may lead even more accurate binding sites prediction. Beyond the realm of binding site prediction, a promising direction for future work is to develop accurate, robust, and scalable computational approaches for protein-DNA or protein-RNA complex structure modeling, capturing protein-DNA and protein-RNA interactions at the atomic level. In this regard, the predicted protein-nucleic acid binding sites can serve as additional restraints, alongside physics- and/or knowledge-guided force fields, to facilitate more efficient and accurate protein-DNA or protein-RNA complex structure modeling. The predicted binding site information can complement and supplement the existing force fields as an additional scoring term to efficiently navigate the conformational space accessible to protein-nucleic acid complexes, leading to improved predictive modeling.

## Data Availability

The raw data used in this study, including the datasets for train, test and validation are collected from publicly available sources and freely available at http://www.csbio.sjtu.edu.cn/bioinf/GraphBind/ and https://github.com/biomed-AI/GraphSite.

## Code Availability

An open-source software implementation of EquiPNAS, licensed under the GNU General Public License v3, is freely available at https://github.com/Bhattacharya-Lab/EquiPNAS.

## Acknowledgements

This work was partially supported by the National Institute of General Medical Sciences (R35GM138146 to D.B.) and the National Science Foundation (DBI2208679 to D.B.).

