## Supplementary Information for "EquiPNAS: improved protein-nucleic acid binding site prediction using protein-language-model-informed equivariant deep graph neural networks"

**Supplementary Table 1.** Hyperparameter optimization using 5-fold cross validation on training dataset for protein-DNA binding sites prediction. Values in bold represent the hyperparameters selected in EquiPNAS for protein-DNA binding site prediction.

| Hyperparameters | Values | 5-Fold Cross Validation ROC-AUC |
|---|---|---|
| Hidden Dimension | 256 | 0.936 |
| | 512 | 0.936 |
| | **768** | 0.938 |
| | 1024 | 0.938 |
| | 1280 | 0.936 |
| Layers | 6 | 0.935 |
| | 8 | 0.937 |
| | 10 | 0.937 |
| | **12** | 0.938 |
| | 14 | 0.937 |
| Dropout Rate | **0.1** | 0.938 |
| | 0.2 | 0.935 |
| | 0.3 | 0.924 |
| $C_\alpha$ - $C_\alpha$ Distance Threshold | 13 | 0.934 |
| | **14** | 0.938 |
| | 15 | 0.936 |

**Supplementary Table 2.** Hyper-parameter optimization using 5-fold cross validation on training dataset for protein-RNA binding sites prediction. Values in bold represent the hyperparameters selected in EquiPNAS for for protein-RNA binding site prediction.

| Hyperparameters | Values | 5-Fold Cross Validation ROC-AUC |
|---|---|---|
| Hidden Dimension | 256 | 0.897 |
| | 512 | 0.9 |
| | **768** | 0.903 |
| | 1024 | 0.902 |
| | 1280 | 0.9 |
| Layers | 6 | 0.897 |
| | 8 | 0.897 |
| | 10 | 0.9 |
| | **12** | 0.903 |
| | 14 | 0.903 |
| Dropout Rate | **0.1** | 0.903 |
| | 0.2 | 0.902 |
| | 0.3 | 0.893 |
| $C_\alpha$ - $C_\alpha$ Distance Threshold | 14 | 0.9 |
| | **15** | 0.903 |
| | 16 | 0.903 |

**Supplementary Table 3.** Protein-DNA binding site prediction performance of EquiPNAS against the competing methods on the Test_129 and Test_181 sets. Values in bold represent the best performance for each input type.

| Input Type | Methods | Test_129 | | Test_181 | |
|---|---|---|---|---|---|
| | | ROC-AUC | PR-AUC | ROC-AUC | PR-AUC |
| *Sequence-only* | SVMnuc* | 0.812 | 0.302 | 0.803 | 0.193 |
| | NCBRPred* | 0.823 | 0.310 | 0.771 | 0.183 |
| | DNAPred* | **0.845** | **0.367** | **0.802** | **0.230** |
| *Using experimental structures* | COACH-D* | 0.710 | 0.269 | 0.655 | 0.172 |
| | NucBind* | 0.811 | 0.294 | 0.796 | 0.191 |
| | DNABind* | 0.858 | 0.402 | 0.825 | 0.219 |
| | GraphBind* | 0.928 | 0.519 | 0.904 | 0.339 |
| | GraphSite | 0.919 | 0.502* | 0.903 | 0.336 |
| | EquiPNAS | **0.943** | **0.582** | **0.921** | **0.393** |
| *Using AlphaFold2 predicted* structural models | COACH-D* | 0.712 | 0.248 | 0.668 | 0.169 |
| | NucBind* | 0.809 | 0.284 | 0.798 | 0.186 |
| | DNABind* | 0.832 | 0.391 | 0.803 | 0.208 |
| | GraphBind* | 0.916 | 0.497 | 0.893 | 0.317 |
| | GraphSite* | 0.934 | 0.544 | 0.917 | 0.369 |
| | EquiPNAS | **0.940** | **0.569** | **0.918** | **0.384** |

Note: * results are obtained directly from the published work of GraphSite.

**Supplementary Table 4.** Protein-RNA binding site prediction performance of EquiPNAS against the competing methods on the Test_117 set. Values in bold represent the best performance for each input type.

| Input Type | Methods | ROC-AUC |
|---|---|---|
| *Sequence-only* | RNABindRPlus* | 0.717 |
| | SVMnuc* | **0.729** |
| | COACH-D* | 0.663 |
| | NucBind* | 0.715 |
| | aaRNA* | 0.771 |
| *Using experimental structures* | NucleicNet* | 0.788 |
| | GraphBind* | 0.854 |
| | EquiPNAS | **0.887** |
| *Using AlphaFold2 predicted structural models* | GraphBind | 0.793 |
| | EquiPNAS | **0.886** |

Note: * results are obtained directly from the published work of GraphBind.

**Supplementary Table 5.** Protein-DNA binding site prediction performance of EquiPNAS using AlphaFold2 predicted structural models as input having moderate confidence predictions (pLDDT ≤ 0.85) and high confidence predictions (pLDDT > 0.85). In all cases, the mean of ROC-AUC and PR-AUC are shown.

| Datasets | All | | pLDDT <=0.85 | | pLDDT > 0.85 | |
|---|---|---|---|---|---|---|
| | Mean PR-AUC | Mean ROC-AUC | Mean PR-AUC | Mean ROC-AUC | Mean PR-AUC | Mean ROC-AUC |
| Test_129 | 0.557 | 0.920 | 0.485 | 0.898 | 0.613 | 0.936 |
| Test_181 | 0.425 | 0.908 | 0.399 | 0.901 | 0.452 | 0.915 |
| Test_117 | 0.378 | 0.867 | 0.311 | 0.855 | 0.450 | 0.879 |

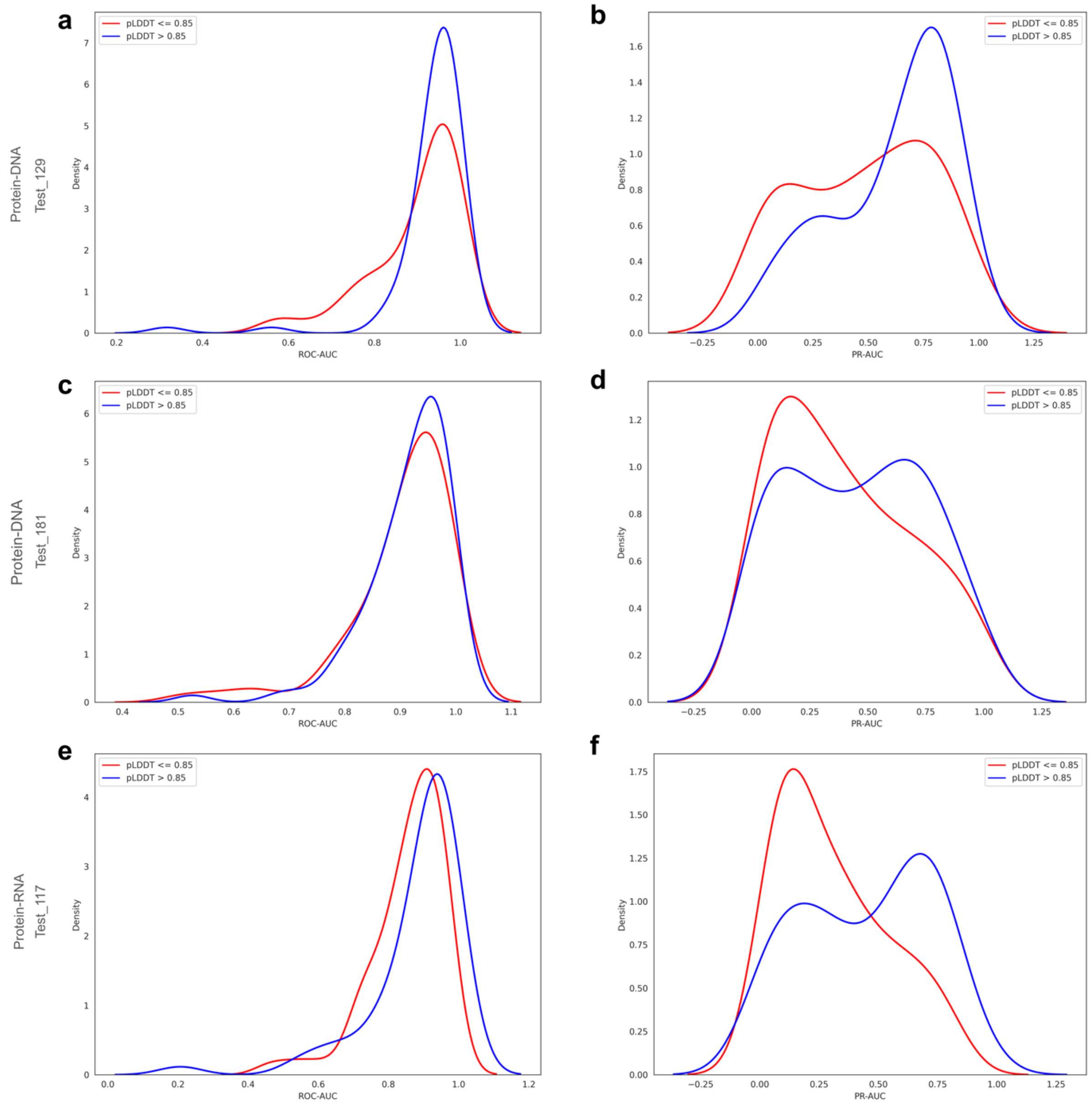


**Supplementary Fig 1.** Accuracy distributions of EquiPNAS protein-nucleic acid binding site prediction using AlphaFold2 predicted structural models as input having moderate confidence predictions (red) and high confidence predictions (blue) in terms of (**a**) ROC-AUC and (**b**) PR-AUC for protein-DNA Test_129; (**c**) ROC-AUC and (**d**) PR-AUC for protein-DNA Test_181; and € ROC-AUC and (**f**) PR-AUC for protein-RNA Test_117.
